# Ventral attention network connectivity is linked to cortical maturation and cognitive ability in childhood

**DOI:** 10.1101/2022.04.12.488101

**Authors:** Hao-Ming Dong, Xi-Han Zhang, Loïc Labache, Shaoshi Zhang, Leon Qi Rong Ooi, B.T. Thomas Yeo, Daniel S. Margulies, Avram J. Holmes, Xi-Nian Zuo

## Abstract

The human brain experiences functional changes through childhood and adolescence, shifting from an organizational framework anchored within sensorimotor and visual regions into one that is balanced through interactions with later-maturing aspects of association cortex. Here, we link this profile of functional reorganization to the development of ventral attention network connectivity across independent datasets. We demonstrate that maturational changes in cortical organization preferentially link to within-network connectivity and heightened degree centrality in the ventral attention network, while connectivity within network-linked vertices predicts cognitive ability. This connectivity is closely associated with maturational refinement of cortical organization. Children with low ventral attention network connectivity exhibit adolescent-like topographical profiles, suggesting that attentional systems may be relevant in understanding how brain functions are refined across development. These data suggest a role for attention networks in supporting age-dependent shifts in cortical organization and cognition across childhood and adolescence.

## Introduction

The human brain undergoes a series of staged developmental cascades across childhood and adolescence progressing from unimodal somatosensory/motor and visual regions through the transmodal association cortex territories that support complex cognitive functions^1–3^. Evidence for the scheduled timing of these neurodevelopmental events has emerged across biological scales, from regional profiles of cellular maturation^4^, synapse formation and dendritic pruning^5^, and intracortical myelination^6^ through macro-scale morphological features including folding patterns^7^ and associated areal expansion^8^. These processes are imbedded within age dependent anatomical changes across lifespan^9^, particularly the prolonged development of association cortex territories. In parallel, substantial progress has been made characterizing the organization^10,11^ and spatiotemporal maturation of large-scale functional systems across the cortex^12,13^. Here, *in vivo* imaging work strongly supports the development of a hierarchical axis, or gradient, of cortical organization, with association territories anchored at the opposite end of a broad functional spectrum from primary sensory and motor regions^14^. Despite clear evidence for age-dependent shifts in the macroscale organization of the cortex from childhood through adolescence, the manner and extent to which specific functional networks may contribute to the widespread process of cortical maturation remains to be determined.

The focused study of discrete functional circuits has provided foundational insights in core maturational processes. For example, discoveries have linked hierarchical changes within amygdala- and ventral striatal-medial prefrontal cortex (mPFC) circuitry to the development of emotional and social functioning in adolescence^1,3,15,16^. Yet, the maturational refinement of these subcortical-cortical circuits does not occur in isolation. Rather, they are embedded within a broad restructuring of functional systems across the cortical sheet^17^. Here, areal and network boundaries become more clearly defined throughout development^18^, while the predominance of local connectivity patterns in childhood gradually gives way to long distance, integrative, connections in adolescence^13,19,20^. This reflects a developmental transition from an anatomically constrained organizational motif to a topographically distributed system^21^. In children, this complex functional architecture is situated within the unimodal cortex, between somatosensory/motor and visual regions. Conversely, adolescents transition into an adult-like gradient, anchored at one end by unimodal regions supporting primary sensory/motor functions and at the other end by the association cortex^14^. While the organizational profiles of large-scale cortical networks are distinct across childhood and adolescence^22^, the extent to which developmental changes within select functional couplings may contribute to the drastic reorganization in the brain hierarchy is an open question. By one view, the developmental transition from unimodal through association cortices reflects the coordinated and shared influence of maturational changes across multiple functional systems spanning the entire connectome. An alternative, although not mutually exclusive, possibility is that specific brain networks may play a preferential role in the widespread developmental refinement of cortical connectivity.

Individual cortical parcels are functionally organized along a global gradient that transitions from somato/motor and visual regions at one end and multimodal association cortex at the other^23^. The hierarchical nature of these functional relationships reflects a core feature of brain organization in both adolescents^14^ and adults^23^. Incoming sensory information undergoes a process of extensive elaboration and attentional modulation as it cascades into deeper layers of cortical processing. Visual system connectivity, as one example, moves along the dorsal and ventral visual streams, uniting within aspects of the dorsal and ventral attention networks including the anterior insula, superior parietal cortex, and operculum parietal before eventually filtering through multimodal convergence zones, particularly within the default network^24^. Although speculative, these data suggest a possible preferential role for sensory orienting and attentional systems in the integrity of the information processing hierarchies in the human brain. Intriguingly, there is mounting evidence to suggest the staged development of a ventral attention network, encompassing aspects of anterior insula, anterior prefrontal cortex and anterior cingulate cortex^11,25^ (see also, cingulo-opercular network^26^ and salience network^27^), that follows the age-dependent shifts in cortical organization across childhood and adolescence^14^. The salience/ventral attentional network, together with frontoparietal network, have been proposed to constitute a dual-network system for the ‘top-down’ and ‘bottom-up’ processing necessary for adaptive behavioral responses^26,28,29^, supporting the functional propagation of information across primary somato/motor, visual, auditory cortex through the default network^24^. These dissociable attentional and control systems are interconnected in children but later segregate over the course of adolescence to eventually form the parallel architectures that support adaptive behavior in adulthood^18^. These data suggest that the attention system may play a preferential role in the transformative brain changes occurring throughout childhood and adolescence. Recent work has also revealed that the lateralization of functional gradients may coincide with attention system lateralization^30^. As such, characterizing the relationships linking attention network connectivity and age-dependent changes in the macroscale brain organization would provide a tremendous opportunity to understand how the functional architecture of cortex is shaped and sculpted across the human lifespan. In turn, this would provide the opportunity to examine how the hierarchical reorganization of the cortical sheet may contribute to the emergence of cognitive and emotional abilities that mark the transition from childhood to adolescence.

In the present study we examined the extent to which specific functional networks may serve to underpin the age-dependent maturation of functional gradient patterns across the cortical sheet. To directly address this open question, we first established the cortical territories exhibiting pronounced functional changes in a longitudinal sample of children and adolescents, revealing the preferential presence of developmental shifts within the ventral attention network. Follow-up analyses excluding regions exhibiting the maximal developmental change in children resulted in the emergence of adolescent-like gradient patterns, suggesting ventral attention territories may play a core role in the expression of adolescent-like connectivity gradients. Moreover, across independent datasets, children with low ventral attention connectivity exhibited a profile of cortical organization that closely resembles prior reports in adolescents and adults. Highlighting the importance of attention network connectivity in cognitive functioning, standardized measures of intelligence linked with reduced attention network degree centrality in children and adolescents. Collectively, these data suggest that ventral attention system functioning in childhood and adolescence may underpin the developmental reorganization and maturation of functional networks across the cortical sheet.

## Results

### Ventral attention network territories demonstrate high degree centrality and pronounced shifts across development

Vertex-level functional connectivity (FC) matrices (20,484×20,484) were first generated using the data provided by the Chinese Color Nest Project (CCNP)^21,32^. In line with prior work^10,11,14^, the top 10% connections of each vertex were retained to enforce sparsity. Degree centrality maps for both children (6 to 12 years of age; n=202) and adolescents (12 to 18 years of age; n=176) were generated to characterize the broad organizational properties of the functional connectome across development (Figure 1A,B). Here, degree centrality reflects the count of above threshold connections for a given vertex to all other vertices (see Methods).

**Figure 1.**
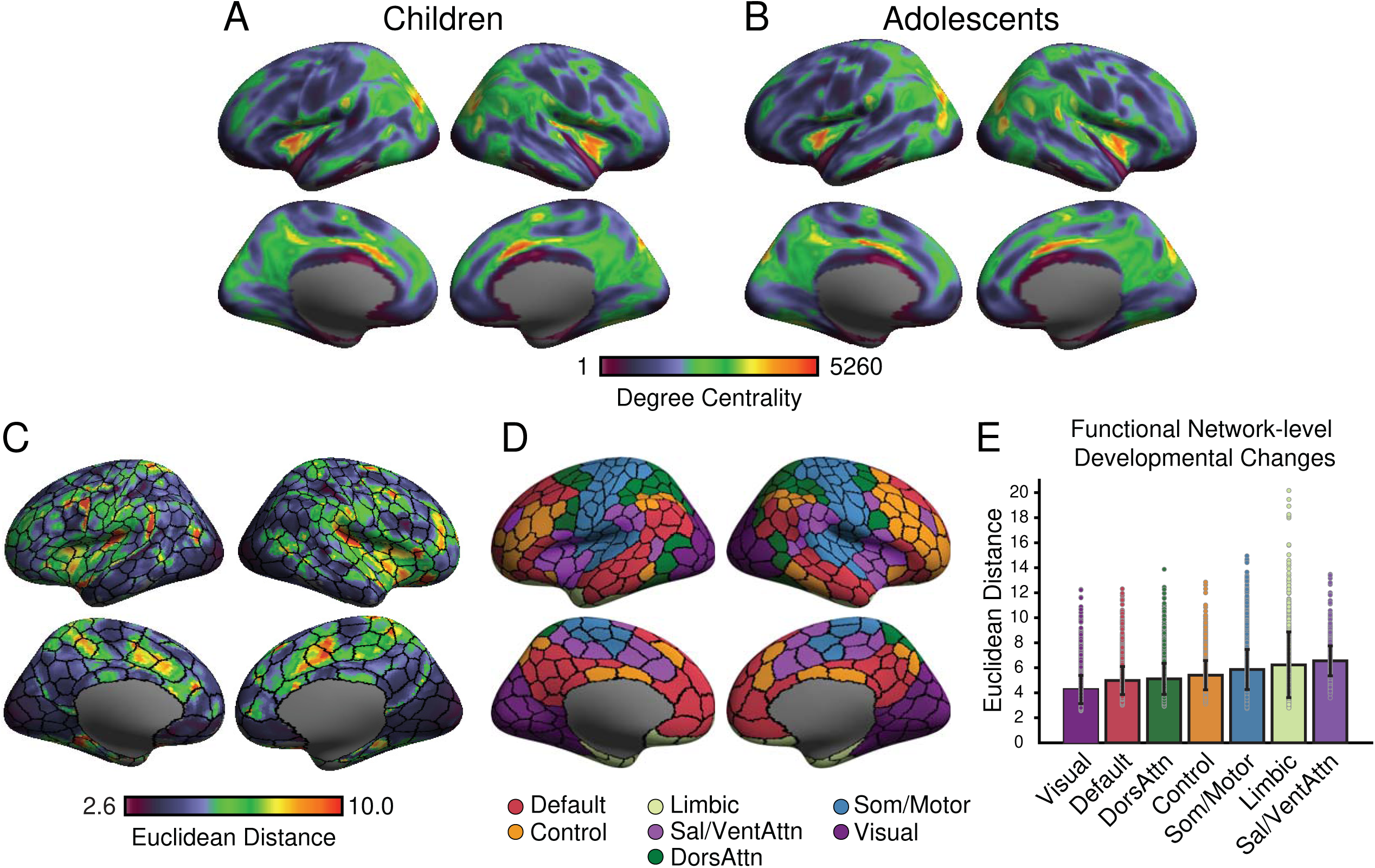
Ventral attention network areas demonstrate high population-level degree centrality but pronounced functional changes across development. Degree Centrality maps in children (A) and adolescents (B) reveal consistent dense connectivity in ventral attention network areas throughout development. Scale bar reflects the count of above threshold connections from a given vertex to all other vertices. Larger values indicate higher degree centrality. (C) Euclidean Distance of the functional connectome at each vertex between children and adolescents reveals a clear switch within the ventral attention network. Larger values indicate greater dissimilarity. (D) Regions based on the Schaefer et al., 400-parcel^31^ atlas and colored by the Yeo et al., 7-network solution^11^. (E) Bar graph reflects changes in the Euclidean Distance of functional connectome at network level (mean network values ± standard error). The ventral attention network (6.5518±1.2002) shows the largest developmental change while visual network (4.2584±1.1257) and default networks (4.9813±1.1207) are most stable between children and adolescents. DorsAttn, dorsal attention (5.1157±1.2499); Sal, salience; Som/Mot, somato/motor (5.8618±1.6067); VentAttn, ventral attention; Control (5.4065±1.1681); and Limbic (6.2337±2.6297).

The observed profiles of degree centrality were highly similar between children and adolescents (Pearson r=0.947, p_spin_≤0.001). Significance was established using permuted spin tests, which preserve the spatial autocorrelation structure of the data^33^. Heightened degree centrality values in both children and adolescents were preferentially evident in aspects of the ventral attention network, including portions of anterior insula, medial prefrontal cortex, and anterior cingulate cortex/midline supplementary motor area (Figure 1). Increased degree centrality was also present in adolescents within default network territories including portions of posterior inferior parietal lobule, posterior cingulate cortex, and precuneus. Additionally, visual system areas including superior and transverse occipital sulcus at the boundary between dorsal and visual network demonstrated high degree centrality values. Conversely, primary somatosensory and motor areas as well as regions within the lateral prefrontal cortex and temporal lobe exhibited relatively low degree centrality. Broadly, these data reflect the presence of dense connectivity within medial and posterior territories along the cortical sheet, while relatively low centrality was evident in lateral prefrontal and somato/motor areas, highlighting a stable pattern of degree centrality across childhood and adolescence.

Degree centrality broadly summarizes profiles of cortical connectivity, to examine developmental changes in functional connectivity strength at the vertex level, we calculated the associated Euclidean distance in functional connectivity similarity between children and adolescents (Figure 1C). Despite the presence of broadly consistent population-level patterns of degree centrality, analyses revealed spatially non-uniform shifts in hub regions of functional connectivity across groups. The maximum developmental changes were anchored within the ventral attention network (Figure 1C-E), encompassing aspects of anterior and posterior insula as well as cingulate cortex. One-way ANOVA revealed the presence of between-network differences (F=790.94, df=6, p≤0.001), with increased Euclidean distance in the ventral attention network, relative to other networks (see multiple comparisons results in Supplemental Table 2). Prior work indicates that maturational age broadly follows the theorized hierarchy of cortical information processing^23^, with somato/motor and visual networks maturing in childhood, while medial prefrontal aspects of default and limbic networks, peak later during adolescence^14,34^. However, in the present analyses, the default network exhibited relatively less developmental change in Euclidean distance between groups, followed by visual network (Figure 1E). Although speculative, these data suggest the presence of specific network-level similarities in connectivity between children and adolescents that may precede broader age-dependent shifts in the macroscale organization of cortex, highlighting the need to consider the manner, in which individual functional networks (e.g., default and attention) are embedded within the broader functional architecture of the brain.

### A core role for the ventral attention network in the macroscale organization of cortex across childhood and adolescence

The transition from childhood to adolescence is marked by pronounced changes in the functional organization of cortex^14,22^. Broadly, this is reflected in the presence of age-dependent transitions across macroscale gradients that extend from unimodal (somato/motor and visual) regions through the cortical association areas that support complex cognition^2,3,15,16^ (Figure 2A). Of note, such a transition is observed not only when restricting analyses to retain the top 10% connections of each vertex to enforce sparsity, but also revealed by when varying sets connections are included in the gradient analyses. Here, the primary transmodal gradient emerges in a higher percentile of excluded connections at each vertex (90%) in adolescents than in children (85%; see (Extended Data Fig.1). Next, we examined whether age-dependent alterations in ventral attention network connectivity might partly account for the maturation of the cortical processing hierarchy as reflected in these overlapping organizing axes, or gradients. Brain areas with maximum differences in Euclidean distance were extracted (See Methods; Figure 2B), and then removed from brain connectivity matrix while we rederived the functional gradients. Here, diffusion map embedding^10,35,36^ was used to decompose participant-level connectivity matrices into a lower dimensional space. The resulting functional components, or gradients, reflect dissociable spatial patterns of cortical connectivity, ordered by the variance explained in the initial functional connectivity matrix^10,14^.

**Figure 2:**
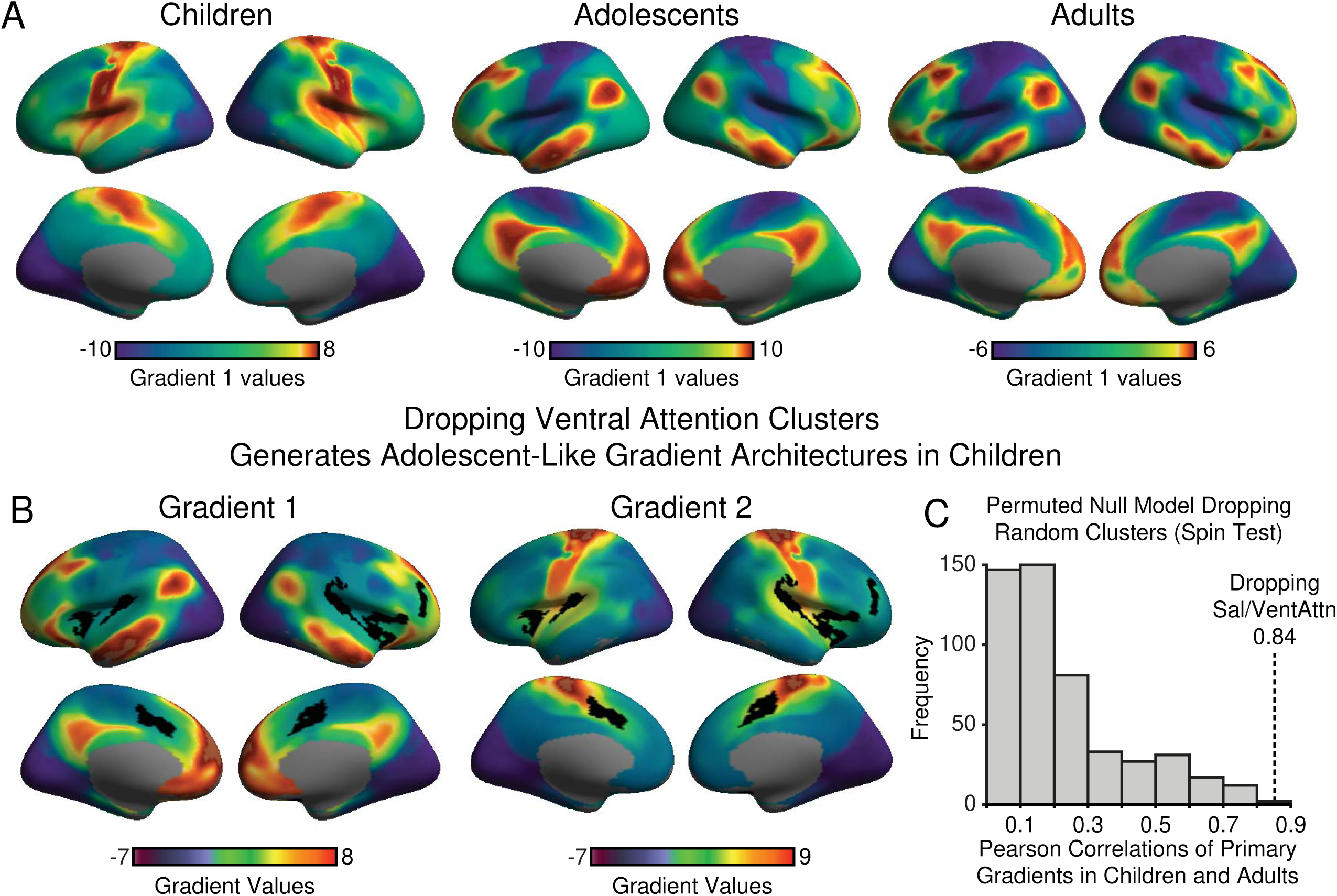
Ventral attention territories play a core role in the expression of adolescent-like connectivity gradients. (A) The principal cortical gradients of connectivity in children, adolescents (data from Dong et al., 2021^14^) and adults (data from Margulies et al., 2016^10^). (B) Clusters with maximum developmental changes in Euclidean Distance were extracted (Figure 1c), denoted as black on the cortical surface. Associated vertices were then dropped from the cortical connectome in the child group prior to rederiving the gradients. Results reveal that default and visual networks anchor the extremes along the principal gradient (Gradient 1), mirroring the principal gradient in both adolescents (r=0.68, p_spin_≤0.001, two-sided spin test) and adults (r=0.84, p_spin_≤0.001, two-sided spin test). In children, the rederived second gradient (Gradient 2) revealed a unimodal architecture separating somato/motor network from visual network, which closely corresponds on the second gradient in both adolescents (r=0.66, p_spin_≤0.001, two-sided spin test) and adults (r=0.85, p_spin_≤0.001, two-sided spin test). (C) To further assess the significance of the results, we constructed a permuted null model in the present data. Clusters with same size and shape as (B) but shuffled locations on the cortical sheet were generated and excluded from analyses in child group 500 times. For each shuffle, the principal gradient was extracted and compared with the principal gradient in adults. The permuted null model shows only 1 case revealing a higher correlation than that observed in the real data, the x-axis indicates the absolute correlation values, the y-axis indicates the frequency of correlations, dotted line refers to the hypothesis being tested.

As identified in our prior study^14^, the primary gradient in children closely matches the second gradient in adolescents and adults. Here, dropping ventral attention areas (a simulation of lesion) generates adolescent- and adult-like gradient architectures in children. The simulated removal of ventral attention network regions led to the formation of a primary gradient in children that closely assembles the first gradient in both adolescents (r=0.68, p_spin_≤0.001) and adults (r=0.84, p_spin_≤0.001). The rederived second gradient in children most closely assembled the second gradient in both adolescents (r=0.66, p_spin_≤0.001) and adults (r=0.85, p_spin_≤0.001). However, while the primary gradient derived from ventral attention network simulated lesioned data in children broadly recapitulated the primary gradient in adults^10^, several inconsistencies were observed. Notably, in the simulated lesioned data from the child group, one end of the primary gradient of connectivity was anchored in the visual areas, with the regions at the other end encompassed broad swaths of the association cortex. Prior work in adults has revealed visual territories along with somato/motor and auditory cortex serve to anchor one end of the primary cortical gradient^10^. Additionally, although the second gradient derived from ventral attention network simulated lesioned data in children closely resembles the second gradient in adults, a muted default network profile can still be observed. While the dropped clusters are not anchored at the extreme end of the primary gradient in children (Figure 2A), it is densely connected and spatially adjacent to somato/motor territories. Further control analysis revealed that areas within ventral attention network primarily contributed to the reversal in gradients. Although speculative, dropping of ventral attention vertices from the gradient analyses may decrease the number of functional connections attributed to somato/motor network, indirectly shifting its position along the gradient spectrum.

Most vertexes from the dropped clusters were from the ventral attention (45.96%) and somato/motor networks (36.57%). Accordingly, the observed transition to an adolescent- and adult-like functional architecture in children may reflect an artifact, resulting from lesioning the data in a manner that removes of aspects of the unimodal territories that anchor the primary gradient in children (Figure 2A). To address this possibility and identify the primary drivers of adolescent-like profiles of brain function in children, we subdivided the dropped clusters into 2 categories: clusters falling within the borders of the somato/motor network and clusters outside the somato/motor network. Gradients were then rederived in the child group to explore the consequences associated with the individual removal of each cluster component. The removal of somato/motor cluster preserves the developmentally typical gradient architecture, in children the first gradient closely resembles the second gradient in adolescents (r=0.94, p_spin_≤0.001) and adults (r=0.91, p_spin_≤ 0.001), while the second gradient closely assemble the first gradient in adolescents (r=0.95, p_spin_≤0.001) and adults (r=0.91, p_spin_≤0.001). Conversely, and consistent with the analyses reported above (Figure 2B), dropping the cluster outside somato/motor network led to the generation of an adolescent- and adult-like gradient architecture in children. Here, the first gradient in children matched the first gradient in adolescents (r=0.78, p_spin_≤0.001), while the second gradient in children resembled the second gradient in adolescents (r=0.78, p_spin_≤0.001). Namely, dropping the clusters outside somato/motor network in children results in a gradient organization that resembles prior reports in adolescents.

We next examined the extent to which the observed elimination of age-dependent shifts in the macroscale organization of the cortex is specific to the removal of ventral attention network-dominated vertices. Here, we generated 500 null models in the child group with clusters dropped at random locations across the cortical sheet, but with shapes and sizes that match the ventral attention network vertices (reflecting the maximum differences in Euclidean Distance between children and adolescents). For each random model, the first and second gradients of child group were extracted and correlated with the corresponding first and second gradients in adults. Providing evidence for the role of the ventral attention network in the formation of adult-like gradient architectures in children, for the primary gradient the observed correlation was greater than the correlations from the null distribution (p≤0.002) across 499/500 permutations (Figure 2C). For the second gradient the observed correlation was greater than the correlations from the null distribution (p≤0.001) across all 500 permutations. While the present analyses are consistent with a core role for the ventral attention network in age-dependent changes in the macroscale organization of the cortex, longitudinal future work should examine the role of person level factors and possible relationships linking large-scale gradient transitions with shifts in attention network functioning across development.

### Cortical development is linked to a pattern of heightened within-ventral attention network connectivity

Extending upon the prior Euclidean Distance analyses, connectome-level changes in functional connectivity between children and adolescents are displayed in a chord diagram (Figure 3) and grouped into networks according to Yeo’s 7-network solution^11^. These data reveal broad increases in within network connectivity for the ventral attention, somato/motor, and default networks as well as a general flattening of cross-network connectivity across development. The within-network connections for the ventral attention, between-network connections for the dorsal attention, and visual networks were increased, while other between-network connectivity with somato/motor, limbic, frontoparietal and default networks are decreased from childhood to adolescence. Individual developmental changes were examined by linear mixed effect (LME) model. Decreased between-network connectivity for the somato/motor network was observed (p≤0.05), while heightened between-network connectivity was evident for the visual (p≤0.05), dorsal attention (p≤0.001), limbic (p≤0.05), and frontoparietal control (p≤0.01) networks. The degree centrality of ventral attention network increased with age in children (p≤0.05), stabilizing in adolescence where no other age-related associations were revealed in within-network connections for ventral attention network.

**Figure 3:**
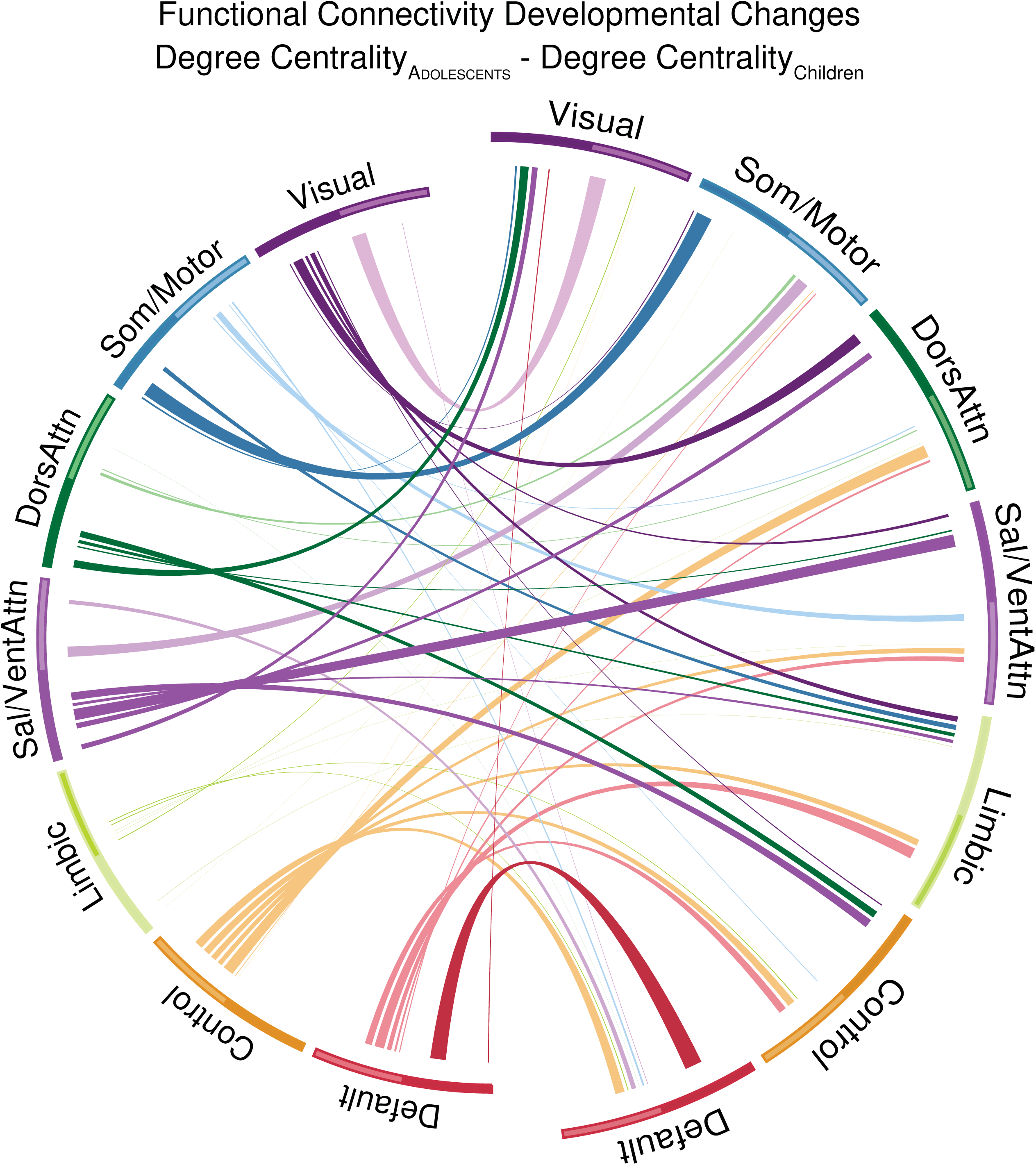
Developmental shifts in functional connectivity between childhood and adolescence. Differences in network-level connectome (The degree centrality matrix of adolescents subtracts the degree centrality matrix of children) functioning between children and adolescents are demonstrated in a chord diagram. For each network, the number connections within the top 10% of each vertex are first summarized at network level, displayed as the links from right half circle with larger radius to left half circle with smaller radius. Reflecting the asymmetrical nature of the thresholded connectivity matrix, links from the left to right half circle indicate the connections within each network that are included in the top 10% of connections to other vertices. Line width highlights the number of connections, with broader lines indicating increased connectivity. Dark colored lines indicate increased values in adolescents, relative to children. Light color lines indicate decrease values in the adolescent group.

When considering the other large-scale association networks, the present analyses suggest a pattern where default network connectivity with other functional systems is pruned during development, indicating a bias towards within network connectivity and an associated differentiation from other network processes. A developmental profile that may coincide with the emergence of the default network at the apex of the network hierarchy in adolescence. Conversely, the frontoparietal control network exhibited increased connectivity with dorsal and ventral attention networks in adolescents, suggesting a potential association between cognitive processes and attentional resources allocation during development.

### Attention network connectivity links with cognitive ability and reveals the presence of adult-like gradient architectures in childhood

The analyses above provide evidence for a relationship between ventral attention network connectivity and the formation of adult-like gradient architectures in children. Significant connectome-level changes were also observed in heightened within-network functional connectivity and degree centrality of ventral attention network (Figure 3). However, it is not yet clear the extent to which individual variability in attention network functioning may link with the adult-like gradient architectures across development. To distinguish it from the typical developmental pattern, we refer to these early-emerging adult-like gradient architectures as an “accelerated maturation pattern”. To examine this potential relationship, we divided participants into subgroups based on their individual ventral attention network connection counts (Figure 4A). Here, to avoid potential bias introduced by confounding factors like scan parameters, population and preprocessing steps, the group split was determined based on the median connection count in the CCNP oldest participants (>17 years of age; dotted line in Figure 4A) instead of independent adult population samples. Participants with fewer connections than this median value were assigned to a low ventral attention group (child n=95 adolescent n=76), all other participants were assigned to a high ventral attention group (child n=107; adolescent n=100). The gradients were then rederived for high and low attention groups across both children and adolescents. No significant associations were observed between the degree centrality of ventral attention network and demographic factors including in age (p=0.45), gender (p=0.82) and head motion (p=0.87). Further comparisons reveal matched demographic features between child the high and low ventral attention groups in age (p=0.0572), gender (p=0.7716) and head motion (p=0.5346).

**Figure 4:**
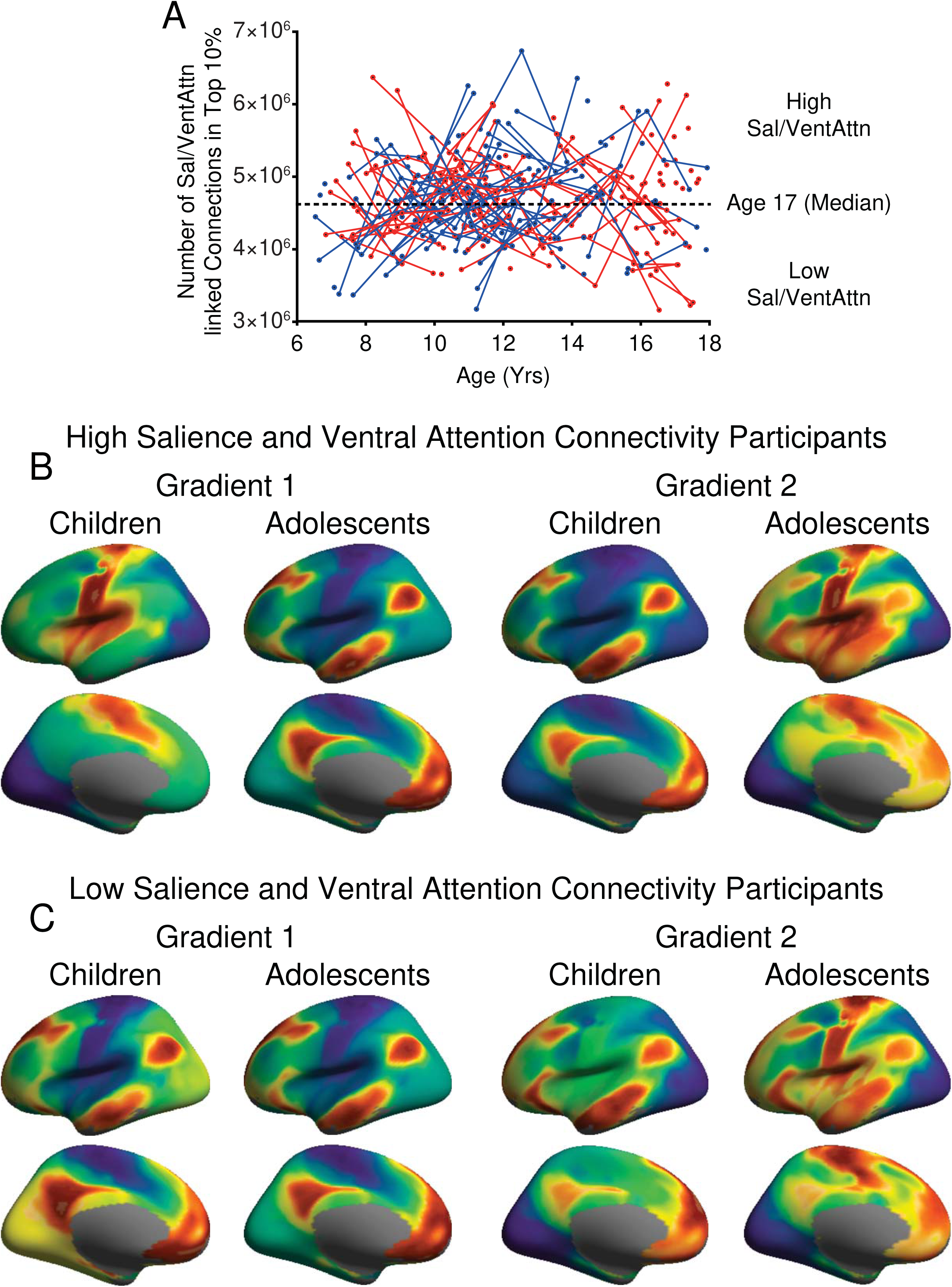
Individual differences in ventral attention network connectivity reveal a functional profile that resembles accelerated cortical maturation in some children. (A) The number of functional connections linked with ventral attention network at individual level. Male participants are marked in blue, female participants are marked in red. Repeated imaging sessions within the same participants are linked by lines. The x-axis represents age range from 6 to 18 years. The y-axis reflects the number of connections that are linked to the ventral attention network following thresholding. Participants were divided into high and low ventral attention connectivity groups according to the median value in age 17-year-old participants (dotted line). (B) The gradient maps in high ventral attention connectivity groups reveal typical patterns identified in our previous work in both children and adolescents^14^. (C) The gradient maps in low ventral attention connectivity groups reveal an accelerated maturation process in children, with both the primary and secondary gradients demonstrated a transmodal architectures. Functional organization was broadly preserved across the high and low attention groups in adolescents, with a subtle muting of the ventral attention gradient values in the low attention participant group. The primary gradient maps in (D) high and low ventral attention connectivity groups derived from longitudinal data. These surface maps display a developmentally normative pattern of gradient reversals in the high ventral attention group children, later scanned in adolescence. Conversely, low ventral attention group children exhibit a stable adolescent-like gradient architecture in both childhood and adolescence. See Extended Data Fig.2 & 3 for the longitudinal analyses of Gradient 1 and Gradient 2 in both hemispheres.

In adolescents, both the primary (r=0.98, p_spin_≤0.001) and secondary (r=0.99, p_spin_≤0.001) gradient architectures were consistent within the low and high ventral attention network participants (Figure 4B,C). For children with high salience and ventral attention connectivity, their functional organization follows the typical pattern of brain maturation^14^ (Figure 2A). Here the primary gradient in the high ventral attention children matched the secondary gradient in both adolescents (r=0.89, p_spin_≤0.001) and adults (r=0.91, p_spin_≤0.001), while the second gradient in high ventral attention children matched the primary gradient in both adolescents (r=0.89, p_spin_≤0.001) and adults (r=0.91607, p_spin_≤0.001). In these children, somato/motor areas were anchored at the opposite extreme from visual regions, revealing a unimodal dominant gradient architecture. However, in children with low ventral attention connectivity we observed a developmentally accelerated pattern of gradient organization that broadly matches the primary and secondary gradients previously identified in both adolescents (Gradient 1: r=0.93, p_spin_≤0.001; Gradient 2: r=0.92, p_spin_≤0.001) and adults (Gradient 1: r=0.68, p_spin_≤0.001; Gradient 2: r=0.65, p_spin_≤0.001).

Permutation analyses were conducted to examine the significance of the dissociable gradient architectures across the groups. Here, 95 and 107 child participants were randomly assigned to two groups corresponding to the number of participants in low and high ventral attention child group for 500 permutations, In the adolescent group, 76 and 100 participants were randomly assigned to two groups corresponding to the number of participants in low and high ventral attention adolescent group for 500 times, then the gradient maps were recomputed and the associated variances were extracted to generate a set of null models.

Children with low ventral attention connectivity exhibit a gradient organization that broadly matches the primary and secondary gradients previously identified in both adolescents and adults. Conversely, children with high ventral attention connectivity exhibit functional organization that follows the typical pattern of brain maturation. The permutation analyses demonstrate that the amount of variance accounted for by the association cortex anchored gradient is increased in the low (Gradient 1 variance: 0.3648) relative to high ventral attention connectivity participants (Gradient 2 variance: 0.1052; p=0, 0/500 permutations revealed greater difference than the real situation). Consistent with this profile the amount of variance accounted for by the unimodal anchored gradient is decreased in the low (Gradient 2 variance: 0.1161) relative to high ventral attention connectivity participants (Gradient 1 variance: 0.3846; p=0, 0/500 revealed greater difference than the real situation).

Conversely, in the adolescence, we did not observe a significant difference between the high and low ventral attention groups in the first association cortex anchored gradient (0.3807 vs. 0.3595, p=0.0620), while the second gradient in high ventral attention adolescent group (0.1052) accounted for a significantly lower amount of variance (p=0.008, 4 in 500 permutations revealed lower difference than the real situation) than low ventral attention adolescent group (0.1161).

The accelerated longitudinal design of the Chinese Color Nest Project^32,37^, which includes longitudinal tracking data with a visiting interval of 1.25 years, provided for additional analyses in participants who were scanned both before (child group) and after their 12th birthday (adolescent group). Here, we identified a set of child participants (n=22) from the low ventral attention group who were also subsequently scanned in their adolescence (n=26, mean age of initial scan=10.98±0.64; mean age of second scan=12.90±0.69). In the participants classified as low ventral attention in childhood, when they transition to adolescence, their first gradient in childhood is highly correlated (r=0.9429, p<0.01) with the first gradient that in their adolescence. A consistent group profile that was also evident when considering their second gradients in both childhood and adolescence (r=0.9353, p<0.01; see Figure 4D).

Conversely, high ventral attention children (n=21), who were also scanned in their adolescence (n=24, mean age of initial scan=11.09±0.63; mean age of second scan=12.94±0.71), displayed a normative developmental trajectory. Here, their first gradient in childhood was highly correlated with their second gradient in adolescence (absolute r=0.9793, p<0.01), while their second gradient in childhood were highly correlated with the first gradient in their adolescence (absolute r=0.9748, p<0.01; see Figure 4D).

If the salience/ventral attention network contributes to the maturation of cortical hierarchy, the functional integrity of attentional systems may also associate with changes in cognitive and behavioral performance. To examine this hypothesis, we next assessed the relationship between standardized measures of cognitive functioning (IQ) and the degree centrality of the ventral attention network with an LME model. LME models for IQ subdomain scores in verbal, perceptual reasoning, working memory, processing speed, and a composite total score were constructed separately. Age, gender, head motion and the vertex-level connectivity counts with ventral attention network were fed into each LME model as fixed effects, repeated measurements were set as the random effect. Results revealed significant negative associations between ventral attention network connectivity and verbal (p≤0.05), perceptual reasoning (p≤0.01) and the composite IQ scores (p≤0.05). Associations with participant age and verbal score (p≤8×10^−6^), perceptual reasoning score (p≤0.05) and total score (p≤0.001), as well as associations with gender and verbal score (p≤0.05), perceptual reasoning score (p≤0.05), and total score (p≤0.05) were also observed. Suggesting that this pattern was not due to variability in data quality, no significant associations between IQ scores and head motion were evident (p>0.05; see LME results in Supplemental Tables 3-7 for each IQ subdomain score).

Collectively, these results demonstrate that individual variability within the attention network in children associates with dissociable motifs of connectivity across the functional connectome and covaries with the cognitive functioning. Children with low ventral attention connectivity exhibit an accelerated profile of cortical maturation that closely resembles prior reports in adolescents and adults. Although additional longitudinal analyses are warranted, these data suggest a fundamental relationship between individual variability within the ventral attention network and the age-dependent changes in the macroscale properties of human brain organization and cognitive ability.

### Ventral attention network connectivity reliably associates with accelerated cortical maturation and cognitive ability across populations

The analyses above reveal a relationship between ventral attention network connectivity, the functional maturation of the cortical connectome, and cognitive functioning in a population of healthy developing children and adolescents from the Chinese Color Nest Project^32^. To examine the generalizability of the above results, we utilized 2186 participants from the ABCD^38^ study, where participant data differed from the CCNP in sample population, study site, MRI scanner, acquisition parameters and longitudinal designs. Here, the ABCD data was limited to participants between 9 to 11 years old. Highlighting the robustness of the analyses reported above, we observed a profile of heightened degree centrality in ventral attention network areas that was consistent with the CCNP analyses (Figure 1A). Broadly, the ABCD degree centrality map demonstrated a lower degree centrality distribution along the cortex than the CCNP dataset (Figure 5A). Here, the inferior parietal gyrus and supramarginal gyrus, together with middle anterior cingulate cortex exhibited highly connected architecture. Hub regions of default network including the angular gyrus, middle temporal gyrus, and post cingulate cortex also revealed heightened degree centrality in the ABCD data, which might again indicate the role of default network as a significant cortical core across the transition to adolescence.

**Figure 5:**
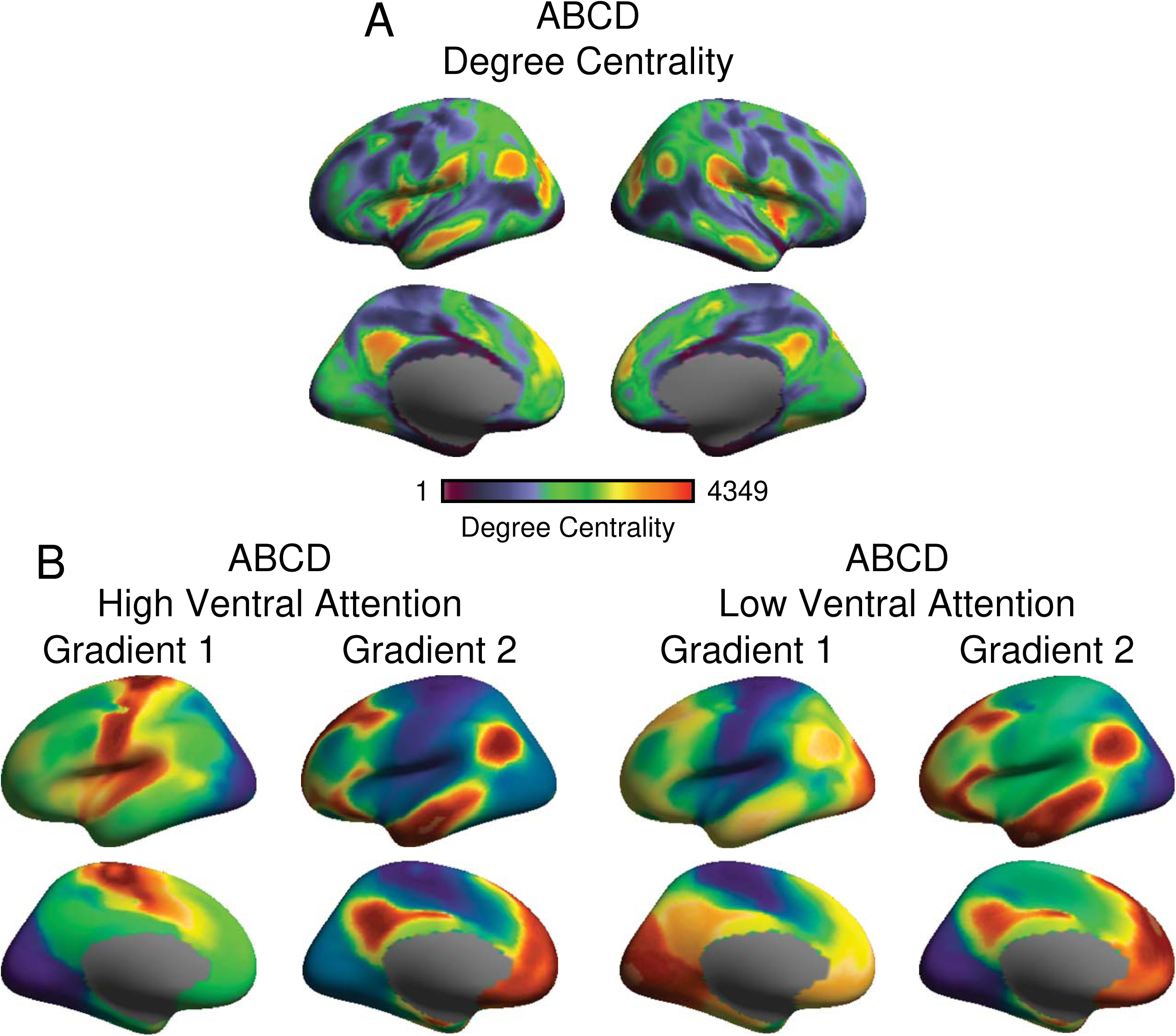
Relationship between ventral attention network connectivity and accelerated cortical maturation reliable across independent datasets. (A) Degree Centrality maps in children (9~11 years old, n=2186) from the ABCD project reveals dense connectivity in ventral attention network areas in a pattern that is consistent with the CCNP participants (Figure 1A). Scale bar reflects the count of above threshold connections from a given vertex to all other vertices. Larger values indicate higher degree centrality. (B) The gradient maps in the ABCD high ventral attention connectivity groups reveal typical patterns identified in our previous work in children^14^ as well as the CCNP high ventral attention group (Figure 4B,C children). (C) The gradient maps in the ABCD low ventral attention connectivity groups closely resemble the profile of accelerated maturation observed in the CCNP sample.

Next, we assessed the reliability of a relationship linking the maturation of functional gradient architecture of the cortical sheet with individual variability in ventral attention connectivity across populations. To do so, we split the children in the ABCD study in a manner consistent with the CCNP analyses to examine whether the ventral attention network connections are coupled with the functional gradient maturation. No significant associations were observed between the degree centrality of ventral attention network and demographic factors like age (p=0.21), gender (p=0.60), head motion (p=0.12) and family income (p=0.15). Further comparisons reveal matched demographic features between child groups in age (p=0.80), gender (p=0.39), head motion (p=0.51) and family income (p=0.15). Consistent with our findings in the CCNP children group, the gradient profiles of children with high ventral attention network connectivity in the ABCD study (n=1367) displayed the developmentally typical gradient profile (Figure 4B). In this group, the primary visual and somato/motor areas occupied the ends of the primary gradient, matched the second gradients in CCNP adolescents (r=0.896, p_spin_≤0.001) and HCP adults (r=0.93, p_spin_≤0.001); while the second gradient revealed transmodal organization matching the first gradients in CCNP adolescents (r=0.918, p_spin_≤0.001) and HCP adults (r=0.944, p_spin_≤0.001). An accelerated developmental profile was revealed in the ABCD children with low ventral attention network links (n=819), broadly matching the primary and secondary gradients previously identified in CCNP adolescents (Gradient 1: r=0.865, p_spin_≤0.001; Gradient 2: r=0.856, p_spin_≤0.001), and demonstrating a hybrid organization comparing with HCP adults (correlations between the first gradient in ABCD low attention group and gradients in HCP adults: r=0.61, p_spin_≤0.001, r=0.73, p_spin_≤0.001; correlations between the second gradient in ABCD low attention group and gradients in HCP adults: r=0.71, p_spin_≤0.001, r=0.62, p_spin_=0.002). These data provide converging evidence, across independent collection efforts, of an association between attention system connectivity and the broader functional organization and maturational properties of cortex.

As a final step we repeated the behavior association analysis in the ABCD project. Here, NIH toolbox scores were used to access the cognitive abilities including crystallized (picture vocabulary and oral reading recognition) and fluid components (pattern comparison processing speed, list sorting working memory, picture sequence memory, flanker test and dimensional change card sort)^39^. As above, a linear regression model was conducted to examine the relationship between the degree centrality of the ventral attention network and cognitive functioning. Age, sex and head motion were set as covariates in the model. Consistent with our CCNP results, these analyses revealed the functional connections of ventral attention network were significantly associated with the cognition total composite standard score (p=0.0024), crystallized composite standard score (p=0.022), cognition fluid composite standard score (p=0.0058), picture vocabulary (p=0.0048), and list sorting working memory (p=0.0025). Other significant associations are reported in the Supplemental Tables. Only the cognition total composite standard score (p=0.0024) and list sorting working memory (p=0.0025) in the ABCD dataset were significant after Bonferroni correction to address multiple comparisons across a total of 15 Linear Mixed Effects (LME) and Linear regression models, no significant association was revealed in the CCNP dataset after the correction.

The replication analysis in the ABCD dataset highlight a potentially key role for the ventral attention network in the maturation process of both cortical hierarchy and cognitive ability. Notably, speaking to the robustness of the observed results, the differences across datasets are quite substantial, including but not limited to participant race, ethnicity, culture, and environment as well as scanner, scanning parameters, socioeconomic factors, and education. In particular, the distinct ethnic/racial composition of study samples can impact the generalizability of brain-behavior associations^40,41^. Here, we applied the connection counts with ventral attention network derived from the CCNP dataset directly on the children in the ABCD dataset, and revealed consistent findings. Across sample collections, children with high ventral attention network connectivity demonstrated typical developmental patterns in functional gradients, while those with lower relative connectivity exhibited adolescent- and adult-liked gradients. The degree centrality of the ventral attention network is significantly associated with similar cognitive components in the CCNP and ABCD datasets. Collectively, these analyses suggest a close and generalizable relationship between the ventral attention network and the process of cortical maturation, as reflected in the presence of macroscale functional gradients. These data are in line with the hypothesis that the ventral attention network may preferentially drive the refinement in the macroscale organization of cortex and cognitive ability during the transition from childhood to adolescence.

## Discussion

A fundamental goal of developmental systems neuroscience is to identify the mechanisms that underlie the scheduled emergence and maturation of the large-scale functional networks central to the information processing capabilities of the human brain. Prior work has demonstrated age-dependent shifts in macroscale organization of cortex^6,9^, revealing the presence of a dominant profile of unimodal cortical organization in children that is later replaced in adolescence by a distributed transmodal architecture spanning from unimodal regions through association cortex. In the current study, we further demonstrated that more connections were required in children to attain a transmodal dominant feature (Extended Data Fig.1), indicating that diffuse connections across broad swaths of association cortex gradually replace the local connections in higher percentiles during adolescence. While this sweeping developmental reorganization of network relationships across the cortical sheet coincides with the transition from childhood to adolescence, reflecting an inflection point at around age 12 to 14, the neurobiological processes underpinning these profound functional changes have yet to be established. Here, by first identifying the densely connected hub regions across development, we demonstrate that ventral attention network regions^11^ encompassing aspects of anterior insula, medial prefrontal cortex, and dorsal anterior cingulate cortex/midline supplementary motor area (also see salience^27^ and cingulo-opercular^26^ networks) may be linked to age-dependent shifts in the macroscale organization of cortex across childhood and adolescence.

Although the attention regions identified in the present analyses are recognized as a single system at a coarse scale^42,43^, this broad architecture is comprised of multiple spatially adjacent but functionally dissociable networks^11,26,27,44^. This complex system is theorized to underpin a cascade of functions supporting information processing, from the control of stimulus driven attention^43^ to the initialization of task sets and maintenance of sustained attention during goal pursuit^29^. Further analysis revealed that virtually lesioning the entire frontoparietal network also generates transmodal organization in the first gradient (See Extended Data Fig.4). Importantly, the salience/ventral attentional network was proposed to have a causal effect on switching between default and frontoparietal networks during task execution^45^. The hierarchical flow of information from sensory regions to deeper levels of cortical processing is also reflected in the spatial continuity of associated functional parcels along cortical surface, for example, the salience network^25,27^, spanning orbital fronto-insular and dorsal anterior cingulate areas through broad posterior areas of insula and dorsal anterior cingulate cortex^25,29^. Broadly, information propagates along posterior to anterior as well as dorsal to ventral axes across cortex and the associated organization of functionally linked parcels is reflected through the presence of continuous functional gradients^10,14^. Here, the ventral attention network is situated at an intermediate position along this functional spectrum transiting from primary sensory/motor networks to the default network that anchors association cortex, which is commonly identified as the first gradient in adolescence and adulthood, while reflecting the second gradient in childhood^14^. This transmodal organizational profile has been inferred to reflect the hierarchy of information flow across cortical territories^10^. Converging evidence for this functional motif has been revealed through analyses of step-wise connectivity. Consistent with classic theories regarding the integration of perceptual modalities into deeper layers of cortical processing^23^, functional relationships spread from primary somato/motor, visual and auditory cortex before converging within ventral attention territories, and eventually the default network^24^. As a whole, these data provide clear evidence situating the ventral attention network between primary unimodal and association cortices, highlighting a role for attention systems in the functional propagation of sensory information to the multimodal regions that support higher-order cognitive functions.

The cingulo-opercular/ventral attention network, together with frontoparietal network, have been proposed to constitute a parallel architecture of executive functioning and cognitive control, supporting adaptive goal pursuit and flexible behavioral adjustments^28^. Although the frontoparietal network, encompassing aspects of dorsolateral prefrontal, dorsomedial prefrontal, lateral parietal, and posterior temporal cortices, is functionally dissociable from cingulo-opercular/ventral attention network in adulthood, studies in developmental populations have revealed that they are functionally linked prior to adolescence^18^. Moreover, in children, the ventral attention network possesses a broad distributed connectivity profile with reduced segregation from salience networks^46^, perhaps reflecting fluid community boundaries in those areas. Anterior prefrontal cortex is shared by the two networks in children but later segregated into ventral attention network in early adolescence, with dorsal anterior cingulate cortex incorporated into frontoparietal network in adulthood^18^. This aspect of network development is consistent with evidence for increased, but more diffuse, patterns of task-evoked activity in children relative to adults^47^, reflecting the presence of scattered pattern of community assignments in prefrontal territories, particularly those supporting attentional processes^22^.

Suggesting a key role for the attentional systems in cognition, our present analyses revealed reliable associations between attention network connectivity and broad measures of intellectual functioning across populations. A recent study also revealed abnormal functioning within limbic and insular networks and associated cognitive deficits in adolescents who were extremely preterm at birth^48^. The maturational course of large-scale brain network coincides with the emergence of adult-like performance on cognitive tasks (e.g., processing speed, shifting and response inhibition)^49,50^. A pattern of attention network development that is theorized to reflect a transition from the over-representation of bottom-up attention to greater top-down attentional and executive functioning capabilites^46^. Converging evidence indicates that ventral attention network functions are intricately linked with the frontoparietal network in both anatomy and function. Although speculative, the corresponding segregation and integration of associated processes likely underpins the maturation of adaptive human cognition.

In the present data, the observed maturational changes were not uniformly distributed across cortical surface, suggesting a key role for the ventral attention network during development. Intriguingly, the greatest functional changes were not evident within the default or primary sensory territories that anchor the functional gradients in adolescence and adulthood. Rather, ventral attention and frontoparietal networks reflected the predominate sources of functional variation across childhood and adolescence (Figure 1E). Of note, developmental changes were not exclusive to attentional network regions, default network and the somato/motor network also demonstrated age related variations across childhood and adolescence. These data are consistent with prior reports suggesting that somato/motor and ventral attention networks are functionally coupled in children, highlighting a distinct community architecture in children relative to adults^22^. Even in adulthood, the ventral areas in somato/motor network can be identified as a separate cluster from dorsal areas as network parcellations increase in their granularity (i.e., 7-network relative to 17-network resolutions in Yeo et al., 2011^11^; or in Gordon et al., 2017^44^).

Prior work has revealed a dramatic developmental change in the topography of ventral attention network, while the spatial organization of default network approaches an adult-like form in childhood^22^. However, it should be noted that the developmental changes in brain’s functional hierarchy are driven by comprehensive reconfigurations across broad swaths of cortex, rather than solely reflecting developmentally mediated shifts in the ventral attention network alone. As illustrated in Figure 3, there are substantial alterations in both between- and within-network connections across different age groups. This is particularly evident in the default network, where a profile of age-linked increases in within-network connectivity replaced between-network connections. These extensive changes along the cortical surface indicate that network organization is not stabilized during development, raising a critical issue that whether a given cortical area belongs to a fixed network across puberty and into adulthood. Future studies should further establish an accurate network affiliation for the developing population. Of note, the size and shape of the human brain undergoes significant changes throughout the lifespan. While it’s indicated that the growth velocities of brain tissues peak before 6 years old^9^, caution is essential when interpreting MRI studies that span development populations. Factors such as head motion and registration errors might bias the results. Therefore, validating with large, independent datasets and alternate methods of assessing brain functioning is crucial to assess the reproducibility, robustness, and generalizability of the current results.

To further characterize how changes in ventral attention network connectivity might underlie the broad functional maturation of cortex, we directly excluded the developmentally dissociable aspects of ventral attention network from the brain connectome and rederived the functional gradients previously characterized in children, adolescence^14^ and adults^10^. The present analyses revealed that virtually lesioning ventral attention clusters characterized by the greatest developmental changes generates adolescent- and adult-like gradient architectures in children, a profile that was specific to attention network territories. Although follow-up work is necessary, our analyses are consistent with the presence of a relative increase in local connections between ventral attention network and adjacent somato/motor territories in childhood, biasing a functional motif in childhood dominated by unimodal organization. Later, diffuse connections across broad swaths of association cortex emerge with age, until eventually the presence of interactions across distinct systems becomes the dominant feature of cortical organization. The increased emphasis on such transmodal integration is consistent with the developmental principle that, over time, local connections are replaced by remote distributed interactions^13^.

Individual level analysis further reveals the influence of ventral attention network on the formation of adult-like functional organization. Children with fewer ventral attention network connections exhibited an accelerated transmodal profile of cortical organization, while the typical unimodal dominated architecture was presented in children with more ventral attention connections. However, an open unanswered question remains the manner through which ventral attention network functioning may reshape the macroscale organization of cortex. Although speculative, the observed results might reflect a consequence of repeated co-activation between ventral attention network regions and associated cortical systems^18^. Accordingly, the dense associations with somato/motor areas^22^ as well as frontoparietal areas^18^ in children indicates ventral attention network is preferentially connected by both primary and association cortex during development, which is also evident in our degree centrality map (Figure 1A and B). While a dense profile of connections may ensure system integrity, excessive or redundant connections are pruned across development in order to optimize efficient information processing as neurons that are not fully integrated within local circuits are eliminated to ensure stable network function^51^. A profile of network segregation and integration that is hypothesized to underpin functional specialization and computational efficiency^19,52^.

Along with the stabilization of its functional organization, cortical maturation is also marked by increased flexibility in resource allocation during task execution. Evidence from task-based fMRI suggests that the salience/ventral attention network, especially the right fronto-insular cortex, plays a critical role in switching between the default mode and frontoparietal networks^45^. This finding supports our hypothesis that the ventral attention network not only serves as a transfer hub between unimodal and transmodal cortices but also coordinates the information flow within the transmodal cortex. In addition to refining the unstable local connections associated with unimodal areas, the ventral attention network may drive the predominance of transmodal organization through a parallel mechanism, which likely involves strengthening the transmodal interactions among higher-order networks to ensure the flexible reallocation of resources. Future work should focus on characterizing the specific properties of ventral attention network across development, for instance, extending to early points in the lifespan, examining its interactions with other network under the background of task execution or considering the roll of individual experience on network development^53^.

Accumulating evidence has revealed a dramatic restructuring of the macroscale organization of the cortex across development, suggesting the scheduled maturation of functional gradient patterns may be critically important for understanding how cognitive and behavioral capabilities are refined across development. However, the underlying developmental processes driving these functional changes remain to be established. Here, by first localizing the significant maturational changes in functional connectome across childhood and adolescence, we demonstrate that the ventral attention network may play a critical role in the onset of the age-dependent shifts that characterize the macroscale organization of cortex across development. Children with fewer functional connections within the ventral attention network exhibit the appearance of an accelerated maturation of gradient architecture and increased cognitive functioning. Although the process of brain maturation emerges through complex interactions across environmental experience and biological systems that span genes and molecules through cells, networks, and behavior, the current findings suggest a core role for attention network-linked territories. The multiscale interactions linking the longitudinal development of the ventral attention network with these maturational processes, and the associated consequences on behavior across health and disease, remains an open question to be answered in future work.

## Acknowledgements

This work was supported by the STI 2030 - the major projects of the Brain Science and Brain-Inspired Intelligence Technology (2021ZD0200500 to X.N.Z.), the National Institute of Mental Health (Grants R01MH120080 and R01MH123245 to A.J.H.), the Major Fund for International Collaboration of National Natural Science Foundation of China (81220108014 to X.N.Z.), the National Basic Science Data Center “Interdisciplinary Brain Database for In vivo Population Imaging” (ID-BRAIN to X.N.Z.) and the Start-up Funds for Leading Talents at Beijing Normal University (to X.N.Z.). BTTY is supported by the NUS Yong Loo Lin School of Medicine (NUHSRO/2020/124/TMR/LOA), the Singapore National Medical Research Council (NMRC) LCG (OFLCG19May-0035), NMRC CTG-IIT (CTGIIT23jan-0001), NMRC STaR (STaR20nov-0003), Singapore Ministry of Health (MOH) Centre Grant (CG21APR1009), the Temasek Foundation (TF2223-IMH-01), and the United States National Institutes of Health (R01MH120080 & R01MH133334). Any opinions, findings and conclusions or recommendations expressed in this material are those of the authors and do not reflect the views of the Singapore NRF, NMRC or MOH.

## Author Contributions

H-MD, AJH and X-NZ designed the research. AJH and X-NZ supervised the research. H-MD and AJH conducted analyses and made figures. X-HZ, LL, S-SZ and LRO conducted validation analyses based on the ABCD dataset. All authors contributed during writing and edited the paper.

## Methods

### Datasets

#### Chinese Color Nest Project (CCNP)

CCNP is a five-year accelerated longitudinal study across the human life span^21,32,54^. A total of 176 scans in adolescents and 202 scans in typically developing children were included in the analysis, details of the dataset and exclusion criteria can be found in our previous work^14^. All MRI data was obtained with a Siemens Trio 3.0T scanner at the Faculty of Psychology, Southwest University in Chongqing. The reported experiments were approved by the Institutional Review Board from Institute of Psychology, Chinese Academy of Sciences. All participants and their parents/guardians provided written informed consent before participating in the study.

#### Adolescent Brain Cognitive Development Study (ABCD)

ABCD is a multi-site longitudinal cohort following the brain and cognition development of over ten thousand 9~10 years old children. MRI scans including T1-weighted, T2-weighted and resting-state fMRI was obtained with 3T Siemens Prisma, General Electric 750 and Phillips scanners across 21 sites, details of the scan parameters can be found in reference^5^. Here, we unitized the MRI baseline data from 2186 children (female 54.4%, mean age 10.01 years old) for the reproducibility analysis. The study was approved by the Institutional Review Board from the University of California, San Diego^55^. All participants and their parents/guardians provided written informed consent^56^.

### MRI Data Preprocessing

#### CCNP dataset

Anatomical T1 images were visually inspected to exclude individuals with substantial head motion and structural abnormalities. Next, T1 images were fed into the volBrain pipeline (http://volbrain.upv.es)^57^ for noise removal, bias correction, intensity normalization and brain extraction. All brain extractions underwent visual inspection to ensure tissue integrity. After initial quality checks, T1 images were passed into the Connectome Computation System (CCS)^58^ for surface-based analyses. CCS pipeline is designed for preprocessing multimodal MRI datasets and integrates publicly available software including SPM^59^, FSL^60^, AFNI^61^ and FreeSurfer^62^. Resting-state fMRI data preprocessing included a series of steps common to intrinsic functional connectivity analyses: (1) dropping the first 10s (4 TRs) for the equilibrium of the magnetic field; (2) estimating head motion parameters and head motion correction; (3) slicing time correction; (4) time series de-spiking; (5) registering functional images to high resolution T1 images using boundary-based registration; (6) removing nuisance factors such as head motion, CSF and white matter signals using ICA-AROMA^63^; (7) removing linear and quadratic trends of the time series; (8) projecting volumetric time series to surface space (the *fsaverage5* model with medial wall masked out); (9) 6mm spatial smoothing. All preprocessing scripts are publicly available on GitHub (https://github.com/zuoxinian/CCS). Any resting-state scan with a mean head motion above 0.5 mm was excluded from further analysis. The demographic information of subjects included in the analyses is listed in Supplemental Table 1.

#### ABCD dataset

Minimally preprocessed T1 images^64^ were fed into Freesurfer^62^ for surface reconstruction. Resting-state fMRI data^64^ preprocessing included a series of steps as following: (1) dropping the initial frames for the equilibrium of the magnetic field; (2) estimating head motion parameters and voxel-wise differentiated signal variance (DVARS) and head motion correction; (3) registering functional images to high resolution T1 images using boundary-based registration; (4) Scrubbing the frames with FD > 0.3 mm or DVARS > 50, along with one volume before and two volumes after. (5) removing nuisance factors such as global signal, head motion, CSF and white matter signals; (6) band-pass filtered (0.009 Hz ≤ f ≤ 0.08 Hz); (7) projecting volumetric time series to surface space (the fsaverage5 model with medial wall masked out); (8) 6mm spatial smoothing. Full details of data preprocessing can be found in previous study^65^. All preprocessing scripts are publicly available (https://github.com/ThomasYeoLab/ABCD_scripts) on GitHub. Any resting-state scan with a max head motion above 5 mm and over half of their volumes censored was excluded from further analysis.

### Degree Centrality Mapping

Functional connectivity (FC) matrices and the corresponding Fisher-z transformed values were first generated for each resting-scan per visit. Then the two test-retest FC fisher-z (FCz) matrices within one visit were averaged to increase signal-to-noise ratio for generating individual FCz matrix for each visit, which was later averaged across individuals to form group-level FCz matrices. For the group-level FCz matrices, only the top 10% functional connenctivities of each vertex were retained, other elements and negative FCs in the matrix were set to 0 to enforce sparsity, yielding an asymmetrical matrix, of which the rows are corresponding to the connectome of each vertex. Degree centrality map was obtained by counting the non-zero elements in each column of the FCz matrix. We then calculated the cosine distance between any two rows of the FCz matrix and subtracted from 1 to obtain a symmetrical similarity matrix, this similarity matrix was later used to derive the gradients.

### Euclidean Distance

To characterize the developmental changes between children and adolescents, the Euclidean distance was computed for each row of the cosine distance matrix between children and adolescents. Then most changed clusters were extracted according to the following two criteria: the top 10% in Euclidean distance map and the cluster size above 500 vertexes. One-way ANOVAs were performed to test the statistical differences between networks.

### Gradients Analysis

The extracted clusters and their FCs were firstly dropped in the initial FCz matrix of children group, and then cosine similarity matrix was calculated. Diffusion map embedding^10,35^ was implemented on the similarity matrix to derive gradients (https://github.com/NeuroanatomyAndConnectivity/gradient_analysis). Within each age bin, the functional connectivity matrix from participants with repeated imaging scans were averaged with scans of other participants to generate a group-level matrix, and then used to derive functional gradients. Pearson correlations were computed between the derived gradients in children and adults. To examine the statistical significance of the observed Pearson correlations, null distributions were generated by randomly rotated the locations of the extracted clusters across the cortical surface while keeping the shape and size fixed. For each permutation, the gradients and correlations were rederived. A total of 500 permutations were performed to generate the null model.

### Chord diagram

Chord diagram was utilized to demonstrate the differences in numbers of functional connectivity between children and adolescents at network level. For each vertex, its connectome was represented as the corresponding row in the FCz matrix. The associated number of network-level connections was obtained by counting the non-0 elements in each network according to the Yeo 7-network solution, generating a matrix with dimension 20,484 (number of vertex) by 7 (number of networks). Vertexes were grouped into networks to generate the final network to network FC matrix (7 by 7). The rows of this network-to-network matrix are displayed as the links from right half circus with larger radius to the left smaller half circus, referring to the FCs with networks in its own connectome, the opposite links from left to right circus represent the columns in the network-to-network matrix, referring to the FCs of each network that were existed in other networks’ connectome (Figure 3).

### High and Low Ventral Attention Network Group Definition

Ventral attention network linked connections were extracted as the corresponding column in network-to-network matrix, referring to the functional connectivity other networks linked with ventral attention network. As we found in prior work^14^, gradient maps became stable in 15-year-old participants, although they still possess a hybrid organization relative to adult participants. The gradient pattern in 17-year-old participants closely resembles what is observed in adults. As such, we hypothesized that the functional connections may reach a stable status at the end of adolescence when compared with younger participants, thus we take the median value of functional connectivity number in 17-year-old age group as the reference. Any single scan with a connectivity number above the threshold was assigned to the “high” ventral attention subgroup, other scans were assigned to the “low” ventral attention group. The gradients were then rederived for each group.

### Association Analysis with IQ Score in the CCNP Dataset

The association between connections number with ventral attention network and IQ scores was estimated with LME model. IQ scores were obtained by Wechsler Children Intelligence Scale IV, including scores in following subdomains: verbal, perception reasoning, working memory, processing speed ability. The LME was conducted using the following formula:

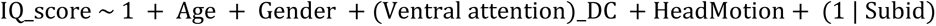

Here (Ventral attention)_DC refers to the connections with ventral attention network, together with age, gender, head motion and the intercept set as the fixed effect factor. Subid refers to the participant IDs, multi measurements for a single participant were coded as an identical nominal variable, set as the random effect factor. LME models were applied for the total IQ and subdomain scores separately.

### Association Analysis with Cognitive Score in the ABCD Dataset

Linear regression model was applied to estimate the association between connections number with ventral attention network and cognitive scores in the ABCD dataset. Cognitive scores were accessed by NIH toolbox, including scores in following domains: crystallized (picture vocabulary and oral reading recognition) and fluid components (pattern comparison processing speed, list sorting working memory, picture sequence memory, flanker test and dimensional change card sort)^10^. The model was conducted using the following formula in MATLAB:

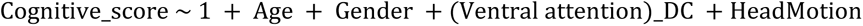

### Developmental Effects on Degree Centrality of Ventral Attention Network

The association between connections number with ventral attention network and age, gender head motion was also estimated with LME model, which was conducted using the following formula in MATLAB:

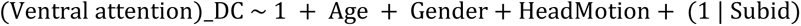

## Data Availability

Data from the CCNP dataset used here are available at Chinese Color Nest Project (CCNP) – Lifespan Brain-Mind Development Data Community at Science Data Bank (https://ccnp.scidb.cn/en) including both anonymized neuroimaging data (https://doi.org/10.57760/sciencedb.07860) and unthresholded whole-brain connectivity matrices grouped by relevant ages (children and adolescents) (https://doi.org/10.11922/sciencedb.00886). The raw CCNP data are available from the website upon reasonable request. The ABCD data used in this report came from the Annual Release 2.0 (https://doi.org/10.15154/1503209) of the ABCD BIDS Community Collection (ABCC; NDA Collection 3165). Source data are provided with this paper.

## Code Availability

Code is available online at our GitHub page: https://github.com/HolmesLab/GradientMaturation

