## Supplementary Text for "Ventral attention network connectivity is linked to cortical maturation and cognitive ability in childhood"

### Supplementary Material

#### The Statistical Tests on Differences in Euclidean Distances

Permutation test was conducted to assess the statistical significance of differences in Euclidean distances between children and adolescents. Participants were randomly divided into two groups, each mirroring the original sample size. Group-level matrices were then created by averaging individual functional connectivity matrices. Cosine similarity was calculated within each group, followed by the extraction of Euclidean distances between two groups. This procedure was repeated 1000 times to generate a null distribution. Cluster-wise significance was examined by averaging the Euclidean Distance within the original identified cluster in the real data and 1000 permutations, the real data has Euclidean Distance (8.67) two times higher value in the real data than the maximum of 1000 permutations (4.24), revealing a significance that exceeds all 1000 permutations ($p=0$ in the current permutation test).

Additional follow-up analyses were conducted at the vertex level. A vertex's Euclidean distance value was considered significantly different only if it exceeded all 1000 permutations ($p=0$ in the current permutation test), ensuring robustness against multiple comparisons. Notably, approximately 80% of the vertices falling within the canonically defined somato/motor and ventral attention networks exhibited significantly greater Euclidean distances than those in the null model. Furthermore, we examined the significance level of the initial cluster reported in the manuscript. 94.21% of this cluster fell within the vertex-level significance map, with the highest effect values observed in the top 10% of Euclidean distances.

**Supplemental Table 1** Demographic composition in the CCNP sample

| Age Range | Total N | Percent Female | FD rest 1 | FD rest 2 |
|---|---|---|---|---|
| 6.0-7.9 | 24 | 50 | 0.213 | 0.119 |
| 8.0-8.9 | 26 | 58 | 0.171 | 0.095 |
| 9.0-9.9 | 36 | 42 | 0.176 | 0.098 |
| 10.0-10.9 | 61 | 59 | 0.162 | 0.089 |
| 11.0-11.9 | 55 | 40 | 0.145 | 0.079 |
| 12.0-12.9 | 39 | 46 | 0.126 | 0.069 |
| 13.0-13.9 | 36 | 58 | 0.112 | 0.061 |
| 14.0-14.9 | 29 | 41 | 0.114 | 0.061 |
| 15.0-15.9 | 22 | 77 | 0.107 | 0.058 |
| 16.0-16.9 | 30 | 77 | 0.090 | 0.050 |
| 17.0-17.9 | 20 | 75 | 0.084 | 0.046 |
| Total Sample | 378 | 54 | 0.139 | 0.077 |
| Children (<12) | 202 | 40 | 0.167 | 0.092 |
| Adolescents (≥12) | 176 | 60 | 0.108 | 0.059 |

Note. FD, framewise displacement, measured in millimeter.

**Supplemental Table 2** Multiple comparisons of Euclidean distance in the ventral attention network relative to other networks

| Network | Network | Differences | p (two-sided) |
|---|---|---|---|
| Ventral Attention | Visual | 2.1797 | <0.001 |
| | SomatoSensory | 0.5834 | <0.001 |
| | Dorsal Attention | 1.3157 | <0.001 |
| | Limbic | 0.4821 | <0.001 |
| | FrontoParietal | 1.263 | <0.001 |
| | Default | 1.6768 | <0.001 |

**Supplemental Table 3** Linear mixed effect models assessing relationships between the degree centrality of the ventral attention network and verbal score

| Fixed effects coefficients | | | | | | |
|---|---|---|---|---|---|---|
| | Estimate | SE | t | DF | p | Adjusted $R^2$ |
| Intercept | 37.573 | 3.4539 | 10.878 | 352 | $5.9102\times10^{-24}$ | 0.82 |
| Age | 0.73539 | 0.16228 | 4.5316 | 352 | $8.0321\times10^{-6}$ | 0.06 |
| Gender | -2.3421 | 0.92919 | -2.5206 | 352 | 0.012156 | 0.03 |
| VA_DC | -0.011269 | 0.0055472 | -2.0315 | 352 | 0.042954 | 0.00918 |
| Head Motion | -0.14552 | 5.6161 | -0.025911 | 352 | 0.97934 | 0 |

**Supplemental Table 4** Linear mixed effect models assessing relationships between the degree centrality of the ventral attention network and perceptual reasoning score

| Fixed effects coefficients | | | | | | |
|---|---|---|---|---|---|---|
| | Estimate | SE | t | DF | p | Adjusted $R^2$ |
| Intercept | 36.688 | 2.8339 | 12.946 | 352 | $1.2746\times10^{-31}$ | 0.8 |
| Age | 0.31133 | 0.13463 | 2.3124 | 352 | 0.02133 | 0.01 |
| Gender | -0.13656 | 0.79135 | -0.17256 | 352 | 0.86309 | 0 |
| VA_DC | -0.011786 | 0.0045036 | -2.617 | 352 | 0.009252 | 0.02 |
| Head Motion | -5.6972 | 4.5735 | -1.2457 | 352 | 0.21371 | 0.00159 |

**Supplemental Table 5** Linear mixed effect models assessing relationships between the degree centrality of the ventral attention network and working memory score

| Fixed effects coefficients | | | | | | |
|---|---|---|---|---|---|---|
| | Estimate | SE | t | DF | p | Adjusted $R^2$ |

| | | | | | | |
|---|---|---|---|---|---|---|
| Intercept | 19.762 | 1.8432 | 10.722 | 352 | $2.124 \times 10^{-23}$ | 0.54 |
| Age | 0.1245 | 0.08727 | 1.4266 | 352 | 0.1546 | 0.00316 |
| Gender | -0.55256 | 0.50879 | -1.086 | 352 | 0.27822 | 0.000858 |
| VA_DC | -0.0023063 | 0.0029386 | -0.78482 | 352 | 0.43309 | 0 |
| Head Motion | -0.27945 | 2.9816 | -0.093724 | 352 | 0.92538 | 0 |

**Supplemental Table 6** Linear mixed effect models assessing relationships between the degree centrality of the ventral attention network and processing speed score

| Fixed effects coefficients | | | | | | |
|---|---|---|---|---|---|---|
| | Estimate | SE | t | DF | p | Adjusted $R^2$ |
| Intercept | 21.957 | 2.284 | 9.6131 | 352 | $1.3994 \times 10^{-19}$ | 0.59 |
| Age | 0.1042 | 0.10953 | 0.95136 | 352 | 0.34207 | 0 |
| Gender | -1.3629 | 0.65933 | -2.0671 | 352 | 0.039454 | 0.02 |
| VA_DC | 0.0019576 | 0.0035967 | 0.54428 | 352 | 0.58659 | 0 |
| Head Motion | -6.652 | 3.6609 | -1.8171 | 352 | 0.070059 | 0.00694 |

**Supplemental Table 7** Linear mixed effect models assessing relationships between the degree centrality of the ventral attention network and composite IQ score

| Fixed effects coefficients | | | | | | |
|---|---|---|---|---|---|---|
| | Estimate | SE | t | DF | p | Adjusted $R^2$ |
| Intercept | 114.07 | 6.3068 | 18.087 | 352 | $3.5824 \times 10^{-52}$ | 0.97 |
| Age | 1.3168 | 0.30704 | 4.2888 | 352 | $2.3237 \times 10^{-5}$ | 0.05 |
| Gender | -4.4193 | 1.9305 | -2.2892 | 352 | 0.022658 | 0.02 |
| VA_DC | -0.021004 | 0.0097763 | -2.1485 | 352 | 0.032358 | 0.01 |
| Head Motion | -10.301 | 9.983 | -1.0318 | 352 | 0.30285 | 0.000139 |

**Supplemental Table 8** Linear regression models assessing relationships between the degree centrality of the ventral attention network and Cognition Total Composite Standard Score (uncorrected) in the ABCD dataset

| | Linear Regression coefficients | | | | |
|---|---|---|---|---|---|
| | Estimate | SE | t | p | Partial $R^2$ |
| Intercept | 0 | 0 | | | |
| Age | 0.3571 | 0.0222 | 16.035 | $8.745\times10^{-55}$ | 0.1040 |
| Gender_Female | 41.766 | 3.363 | 12.42 | 0 | 0 |
| Gender_Male | 41.678 | 3.380 | 12.33 | 0 | |
| VA_DC | 0.01218 | 0.004 | 3.04 | 0.0024 | 0.0037 |
| Head Motion | -23.89 | 4.6897 | -5.0937 | $3.8\times10^{-7}$ | 0.0105 |

**Supplemental Table 9** Linear regression models assessing relationships between the degree centrality of the ventral attention network and Crystallized Composite Standard Score (uncorrected) in the ABCD dataset

| | Linear Regression coefficients | | | | |
|---|---|---|---|---|---|
| | Estimate | SE | t | p | Partial $R^2$ |
| Intercept | 0 | 0 | | | |
| Age | 0.2638 | 0.0185 | 14.291 | 0 | 0.0849 |
| Gender_Female | 53.371 | 2.787 | 19.149 | 0 | 0.0017 |
| Gender_Male | 53.928 | 2.80 | 19.251 | 0 | |
| VA_DC | 0.0076 | 0.0033 | 2.2925 | 0.022 | 0.0022 |
| Head Motion | -13.28 | 3.886 | -3.416 | $6.45\times10^{-4}$ | 0.0049 |

**Supplemental Table 10** Linear regression models assessing relationships between the degree centrality of the ventral attention network and Cognition Fluid Composite Standard Score (uncorrected) in the ABCD dataset

| | Linear Regression coefficients | | | | |
|---|---|---|---|---|---|
| | Estimate | SE | t | p | Partial $R^2$ |
| Intercept | 0 | 0 | | | |
| Age | 0.337 | 0.0265 | 12.74 | 0 | 0.0682 |
| Gender_Female | 49.6 | 4 | 12.4 | 0 | 0.0013 |
| Gender_Male | 48.89 | 4 | 12.16 | 0 | |
| VA_DC | 0.013 | 0.0048 | 2.7614 | 0.0058 | 0.0032 |
| Head Motion | -27 | 5.579 | -4.84 | $1.38\times10^{-6}$ | 0.0099 |

**Supplemental Table 11** Linear regression models assessing relationships between the degree centrality of the ventral attention network and Picture Vocabulary Standard Score (uncorrected) in the ABCD dataset

| | Linear Regression coefficients | | | | |
|---|---|---|---|---|---|
| | Estimate | SE | t | p | Partial $R^2$ |
| Intercept | 0 | 0 | | | |
| Age | 0.29 | 0.02 | 13.34 | 0 | 0.0748 |
| Gender_Female | 47.76 | 3.28 | 14.57 | 0 | 0.0029 |
| Gender_Male | 48.61 | 3.30 | 14.75 | 0 | |
| VA_DC | 0.0086 | 0.0039 | 2.21 | 0.027 | 0.0021 |
| Head Motion | -12.9 | 4.57 | -2.83 | 0.0048 | 0.0034 |

**Supplemental Table 12** Linear regression models assessing relationships between the degree centrality of the ventral attention network and Pattern Comparison Processing Speed Standard Score (uncorrected) in the ABCD dataset

| | Linear Regression coefficients | | | | |
|---|---|---|---|---|---|
| | Estimate | SE | t | p | Partial $R^2$ |
| Intercept | 0 | 0 | | | |
| Age | 0.41 | 0.039 | 10.33 | 0 | 0.0459 |
| Gender_Female | 42.32 | 5.91 | 7.16 | $1.14\times10^{-12}$ | 0.0064 |
| Gender_Male | 40.06 | 5.95 | 6.74 | $2.05\times10^{-11}$ | |
| VA_DC | 0.006 | 0.007 | 0.86 | 0.39 | 0.0003 |
| Head Motion | -36.18 | 8.25 | -4.39 | $1.2\times10^{-5}$ | 0.0083 |

**Supplemental Table 13** Linear regression models assessing relationships between the degree centrality of the ventral attention network and Flanker Inhibitory Control and Attention Standard Score (uncorrected) in the ABCD dataset

| | Linear Regression coefficients | | | | |
|---|---|---|---|---|---|
| | Estimate | SE | t | p | Partial $R^2$ |
| Intercept | 0 | 0 | | | |
| Age | 0.19 | 0.023 | 8.16 | $5.6\times10^{-16}$ | 0.0295 |
| Gender_Female | 67.22 | 3.51 | 19.16 | 0 | 0.0020 |
| Gender_Male | 67.96 | 3.53 | 19.27 | 0 | |
| VA_DC | 0.011 | 0.0042 | 2.67 | 0.23 | 0.0031 |
| Head Motion | -5.8 | 4.89 | -1.18 | 0.24 | 0.0006 |

**Supplemental Table 14** Linear regression models assessing relationships between the degree centrality of the ventral attention network and List Sorting Working Memory Standard Score (uncorrected) in the ABCD dataset

| | Linear Regression coefficients | | | | |
|---|---|---|---|---|---|
| | Estimate | SE | t | p | Partial $R^2$ |
| Intercept | 0 | 0 | | | |
| Age | 0.2 | 0.03 | 6.4 | $1.86\times10^{-10}$ | 0.0183 |
| Gender_Female | 67.56 | 4.74 | 14.25 | 0 | 0.0005 |
| Gender_Male | 68.04 | 4.76 | 14.28 | 0 | |
| VA_DC | 0.017 | 0.006 | 3.02 | 0.0025 | 0.0041 |
| Head Motion | -18.29 | 6.61 | -2.77 | 0.0057 | 0.0034 |

**Supplemental Table 15** Linear regression models assessing relationships between the degree centrality of the ventral attention network and Dimensional Change Card Sort Standard Score (uncorrected) in the ABCD dataset

| | Linear Regression coefficients | | | | |
|---|---|---|---|---|---|
| | Estimate | SE | t | p | Partial $R^2$ |
| Intercept | 0 | 0 | | | |
| Age | 0.26 | 0.024 | 10.87 | 0 | 0.0510 |
| Gender_Female | 62.45 | 3.57 | 17.51 | 0 | 0.0017 |
| Gender_Male | 61.75 | 3.58 | 17.23 | 0 | |
| VA_DC | 0.006 | 0.004 | 1.33 | 0.18 | 0.0008 |
| Head Motion | -16.7 | 4.97 | -3.36 | $8\times10^{-4}$ | 0.0049 |

**Supplemental Table 16** Linear regression models assessing relationships between the degree centrality of the ventral attention network and Picture Sequence Memory Standard Score (uncorrected) in the ABCD dataset

| Linear Regression coefficients | | | | | |
|---|---|---|---|---|---|
| | Estimate | SE | t | p | Partial $R^2$ |
| Intercept | 0 | 0 | | | |
| Age | 0.16 | 0.034 | 4.62 | $4\times10^{-6}$ | 0.0096 |
| Gender_Female | 84 | 5.10 | 16.47 | 0 | 0.0011 |
| Gender_Male | 83.24 | 5.13 | 16.23 | 0 | |
| VA_DC | 0.008 | 0.006 | 1.29 | 0.196 | 0.0008 |
| Head Motion | -19.31 | 7.12 | -2.71 | 0.007 | 0.0033 |

**Supplemental Table 17** Linear regression models assessing relationships between the degree centrality of the ventral attention network and Oral Reading Recognition Standard Score (uncorrected) in the ABCD dataset

| Linear Regression coefficients | | | | | |
|---|---|---|---|---|---|
| | Estimate | SE | t | p | Partial $R^2$ |
| Intercept | 0 | 0 | | | |
| Age | 0.21 | 0.019 | 11.048 | 0 | 0.0527 |
| Gender_Female | 66 | 2.8 | 23.61 | 0 | 0.0002 |
| Gender_Male | 66.23 | 2.81 | 23.56 | 0 | |
| VA_DC | 0.0051 | 0.003 | 1.52 | 0.13 | 0.0010 |
| Head Motion | -12.55 | 3.9 | -3.22 | 0.0013 | 0.0045 |
